# Structure-based discovery of small molecule inhibitors of the autocatalytic proliferation of α-synuclein aggregates

**DOI:** 10.1101/2021.12.05.471256

**Authors:** Sean Chia, Z. Faidon Brotzakis, Robert I. Horne, Andrea Possenti, Benedetta Mannini, Rodrigo Cataldi, Magdalena Nowinska, Roxine Staats, Sara Linse, Tuomas P. J. Knowles, Johnny Habchi, Michele Vendruscolo

**Affiliations:** Centre for Misfolding Diseases, Department of Chemistry, University of Cambridge, Cambridge CB2 1EW, UK; Department of Biochemistry & Structural Biology, Center for Molecular Protein Science, Lund University, 221 00 Lund, Sweden; Department of Physics, Cavendish Laboratory, Cambridge CB3 0HE, UK

**Keywords:** Parkinson’s disease, α-synuclein, protein aggregation, computational docking, structure-based drug discovery, kinetic-based drug discovery

## Abstract

The presence of amyloid fibrils of α-synuclein is closely associated with Parkinson’s disease and related synucleinopathies. It is still very challenging, however, to systematically discover small molecules that prevent the formation of these aberrant aggregates. Here, we describe a structure-based approach to identify small molecules that specifically inhibit the surface-catalyzed secondary nucleation step in the aggregation of α-synuclein by binding to the surface of the amyloid fibrils. The resulting small molecules are screened using a combination of kinetic and thermodynamic assays for their ability to bind α-synuclein fibrils and prevent the further generation of toxic oligomers. This study demonstrates that the combination of structure-based and kinetic-based drug discovery methods can lead to the identification of small molecules that selectively inhibit the autocatalytic proliferation of α-synuclein aggregates.

## Introduction

Parkinson’s disease is the most common neurogenerative movement disorder, which affects over 6 million individuals worldwide (1–4). This disease is characterised histopathologically by the accumulation of aberrant deposits known as Lewy bodies, which are composed primarily of the aggregated form of the intrinsically disordered protein α-synuclein (5, 6). Aggregates of α-synuclein, including soluble misfolded oligomers and insoluble amyloid fibrils, can induce neurotoxicity through a multitude of mechanisms, including cell membrane disruption and mitochondrial damage, which ultimately cause neuronal death (7–9).

Because of the relevance α-synuclein aggregates, much effort has been devoted towards the characterization of their structures (10–14). These structures have enabled the development of structure-based drug discovery approaches, including in particular the identification of peptide-based inhibitors to prevent α-synuclein aggregation (15, 16). Furthermore, binding sites along the surface of α-synuclein fibrils for the development of diagnostics tools have also been identified (17–20). These developments are relevant considering the current lack of radiotracers for measuring the accumulation of α-synuclein aggregates in the human brain, and that such diagnostics tools may eventually enable the presymptomatic diagnosis of synucleinopathies (17, 21–23).

Previous studies have used high-throughput docking approaches to identifying α-synuclein fibril-binding compounds (17–20). However, because of the great technical difficulties in establishing reproducible high-throughput kinetic assays to monitor α-synuclein aggregation, the experimental validation of the compounds predicted from computational screenings has been challenging. Recent advances in chemical kinetics approaches have allowed the identification of small molecules and molecular chaperones that are able to inhibit α-synuclein aggregation (24–27). It has thus been possible to inhibit specifically the surface-catalysed secondary nucleation step, which is responsible for the autocatalytic proliferation of α-synuclein fibrils, by binding competitively with α-synuclein monomers along specific sites on the surface of α-synuclein fibrils (24, 26). This kinetic-based approach is particularly advantageous as it allows experiments to be performed in a high-throughput manner in 96-well plates, while conferring quantitative analysis of the effect of the compounds on specific microscopic steps in the aggregation process of α-synuclein. These quantitative measurements subsequently enable structure-activity relationship (SAR) studies and facilitate the systematic optimization of the compounds properties (25, 28).

In the present study, we have identified small molecules that bind to α-synuclein fibrils in order to make advances on two problems: (1) how to prevent the fibril-catalyzed secondary nucleation in the autocatalytic proliferation of α-synuclein fibrils, and (2) how to identify molecular tracers for measuring the accumulation of α-synuclein aggregates through imaging methods. Towards these goals, we demonstrate the use of an *in silico* and *in vitro* combinatorial framework in identifying compounds that bind to α-synuclein fibrils, and subsequently inhibit the secondary nucleation process in the aggregation of α-synuclein. In this framework, we first employ a computational method that combines two docking techniques to identify compounds with high predicted binding affinity for α-synuclein fibrils. This list is then validated experimentally through chemical kinetics, which identifies the top compounds that are able to inhibit the surface-catalysed secondary nucleation step in the aggregation of α-synuclein. The binding affinity of these compounds for α-synuclein fibrils is then validated experimentally.

Overall, this strategy demonstrates the rational development of a combined structure-based and kinetic-based framework to identify compounds that can bind to α-synuclein fibrils, and presents an opportunity for the systematic development of small molecules that can be potentially used as therapeutic and diagnostic tools for synucleinopathies.

## Results

### A framework to identify small molecules that bind α-synuclein fibrils

In this work we describe a framework to identify small molecules that bind specific sites on the surface of α-synuclein fibrils and are able to block the process of fibril-catalysed secondary nucleation. This method consists of a computational docking approach to identify small molecule candidates from a large library of compounds, and a subsequent *in vitro* approach based on chemical kinetics to assess the ability of the candidates to inhibit the aggregation of α-synuclein, as well as their affinity towards α-synuclein fibrils (**Fig. 1**).

**Figure 1.**
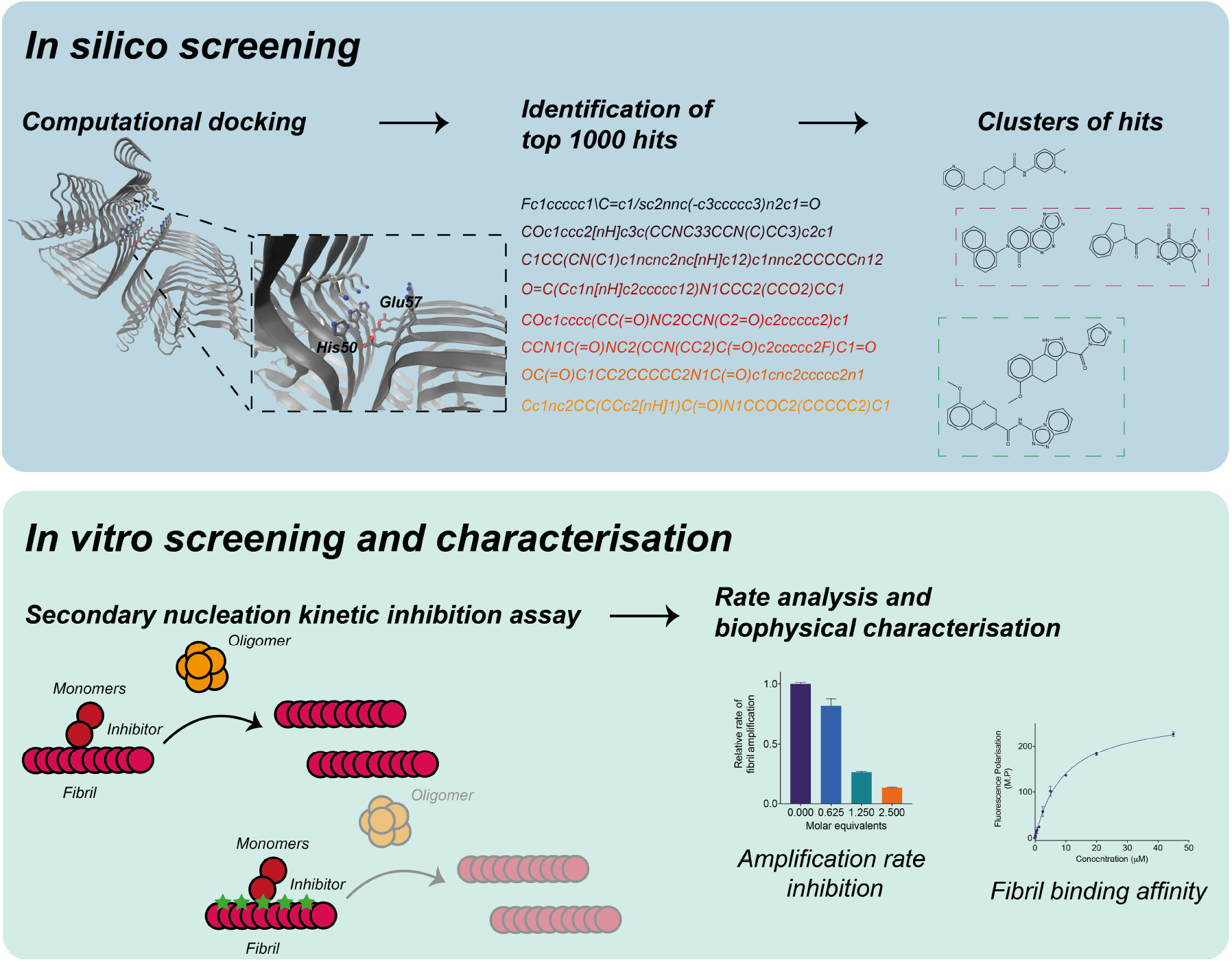
A combined structure-based and kinetic-based approach to identify small molecules that bind α-synuclein fibrils and inhibit its aggregation. In the first step, computational docking is performed on a large library of small molecules. The top candidates are then clustered to identify a subset of chemically diverse molecules that exhibit high predicted binding scores for μ-synuclein fibrils. Subsequently, these small molecules are experimentally validated through a kinetic assay for their ability to inhibit the secondary nucleation aggregation of α-synuclein by binding to the surface of fibrils. Further rate constant analysis and fibril binding experiments allows for the positive molecules to be characterised based on both their inhibition of the kinetic assay, as well as their binding affinity towards α-synuclein fibrils.

Firstly, from a library of compounds, small molecules are individually docked against a binding pocket chosen along the groove of the fibril (Materials and Methods). This groove, which involves residues His50 and Glu57, was selected as the potential site for docking since its geometry and position identified it as a likely catalytic site for α-synuclein secondary nucleation (**Fig. 1**). Using the predicted binding scores (ΔG_b_) as the parameter for ranking the compounds, the docking procedure generated a list of top 1,000 candidates. As a means of increasing the chemical diversity of the compounds to be experimentally validated, a clustering method based on the chemical similarity of the compounds was adopted, in which the centroids of each of the clusters were selected as the final candidates of the library of compounds to be tested *in vitro* (**Fig. 1**).

Next, these centroid compounds were validated experimentally using a chemical kinetics-based assay of α-synuclein aggregation. Specifically, in the presence of low amounts of pre-formed seeds and at acidic pH, the aggregation of α-synuclein is dominated by a surface-catalysed secondary nucleation process, in which monomers form nuclei along the fibril surfaces (29). This autocatalytic process results in the rapid generation of aggregates that then elongate to form α-synuclein fibrils. When the aggregation of α-synuclein is performed in the presence of small molecules with suitable binding affinity for the surface of the α-synuclein fibrils, this results in a decreased amplification rate of α-synuclein aggregates (24, 26).

The small molecule inhibitors identified in this way were further validated for their binding affinity towards α-synuclein fibrils using fluorescence polarisation and mass-spectrometry based pull-down assays. Overall, using this framework, candidate small molecules could be characterised through their predicted and experimental binding affinity, as well as their kinetic inhibitory properties as a result of their interaction with α-synuclein fibrils.

### In-silico docking of compounds predicted to bind α-synuclein fibrils

Using AutoDock Vina (30) and FRED (31), a wide distribution of binding scores was observed for compounds in the ZINC library (32) (Materials and Methods). When assessing the top 10,000 compounds with the highest binding affinity, we observed a wider distribution of binding scores using FRED (−4 to −14 kcal/mol) than AutoDock Vina (−6.6 to −8.6 kcal/mol) (**Fig. S1**). We also found that the predicted binding scores did not correlate strongly between the two methods (**Fig. S1**). Such differences in the predicted scores can be attributed to the different scoring functions used among docking methods, leading to the choice of consensus compounds (33). Thus, we selected the top 10% (1,000 common compounds) of the candidates with a high predicted binding score in both methods (**Fig. S1**). The candidate library was further refined by employing a clustering method based on the chemical structures of the compounds to identify the centroids of each cluster as the representative molecule in the cluster. Although this procedure may result in a lower number of hits due to the exclusion of chemical derivatives of a potential α-synuclein fibril binder that are clustered together, it also increases the chemical diversity of the library set, and therefore allows the sampling of a wider chemical space to screen for potential fibril binders.

### Identification of compounds that inhibit α-synuclein secondary nucleation

From the library of centroids, we selected 67 compounds for experimental validation in terms of their binding affinity towards α-synuclein fibrils (Materials & Methods). Out of the 67 molecules tested, 5 molecules were found to inhibit α-synuclein aggregation (**Fig. 2**). The small molecules were found to inhibit the aggregation of α-synuclein to different extents. In particular, 3 of the molecules (A, B, and E) showed moderate potency, increasing the half-time (*t*_*1/2*_) of the aggregation by 1.5 times, while molecules C and D exhibited stronger potency by increasing the *t*_*1/2*_ of the aggregation by 3.5 and 3 times, respectively. We also found that the chemical structures of these inhibitors tend to involve aromatic moieties (**Fig. 2C**). More specifically, the aromatic regions of these molecules appeared within close proximity to the residues along the groove of the α-synuclein fibrils, suggesting that interactions could be established between the small molecules and α-synuclein through these regions (**Fig. 3**). Furthermore, we also observed a high similarity in terms of the small molecule positions within the selected groove of α-synuclein fibrils between using docking methods, thus suggesting that the binding of these small molecules to the α-synuclein fibrils may involve specific interactions, which are likely to be a combination of electrostatic and non-polar nature (**Fig. 3**).

**Figure 2.**
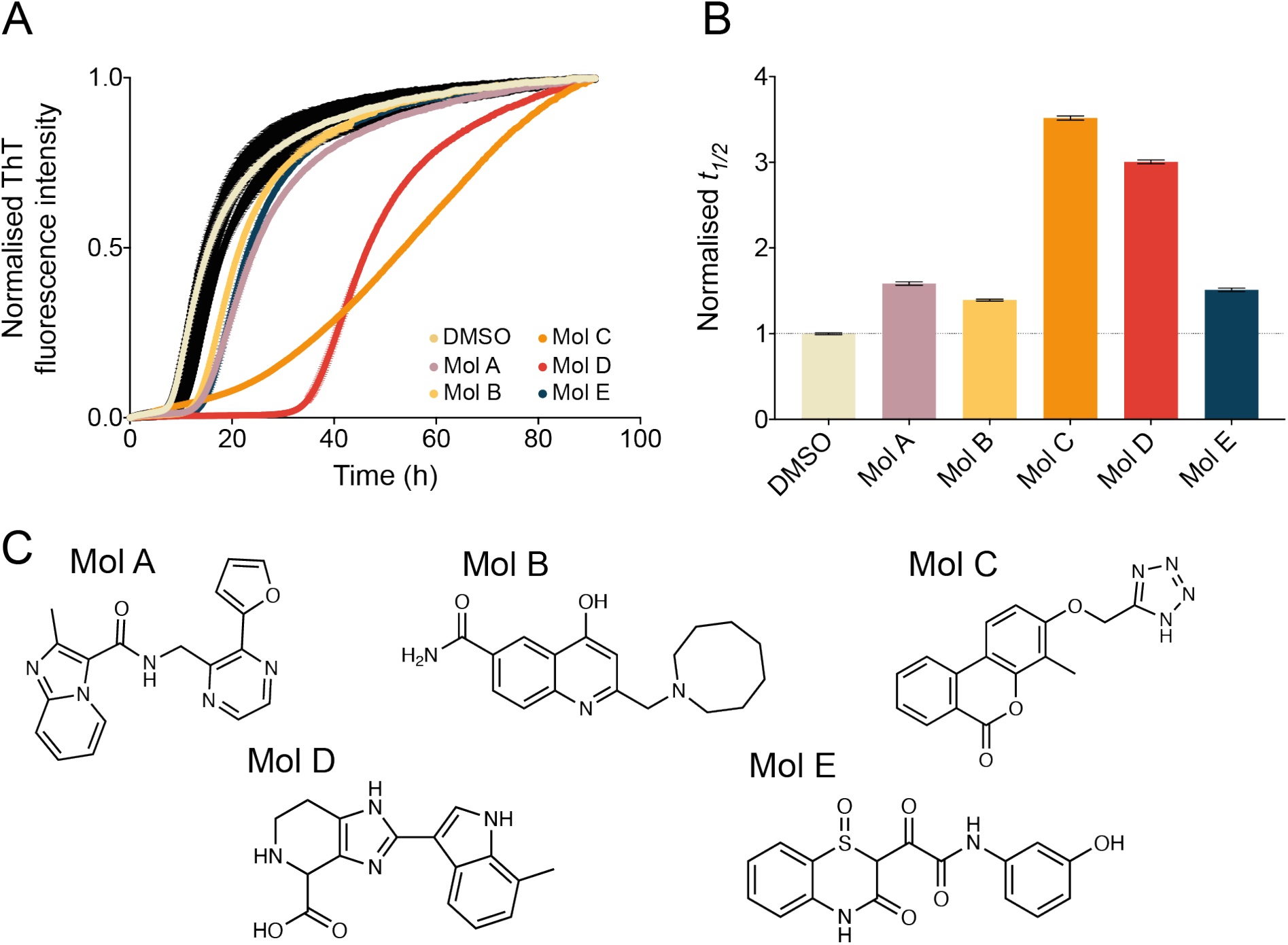
Molecules identified from the docking library inhibit the aggregation of α-synuclein. **(A)** Kinetic profiles of a 10 μM solution of α-synuclein in the presence of 25 nM seeds at pH 4.8, 37 °C, either in the presence of 1% DMSO alone (beige), in the presence of 10 molar equivalents of molecules A-E (represented in different colours), or in the presence of 10 molar equivalents of other representative molecules in the docking library (black). **(B)** Relative *t*_*1/2*_ of the aggregation of α-synuclein in the presence of molecules A-E as shown in panel A, normalised to the DMSO control. **(C)** Chemical structures of the molecules A-E, which are able to significantly inhibit the aggregation of α-synuclein.

**Figure 3.**
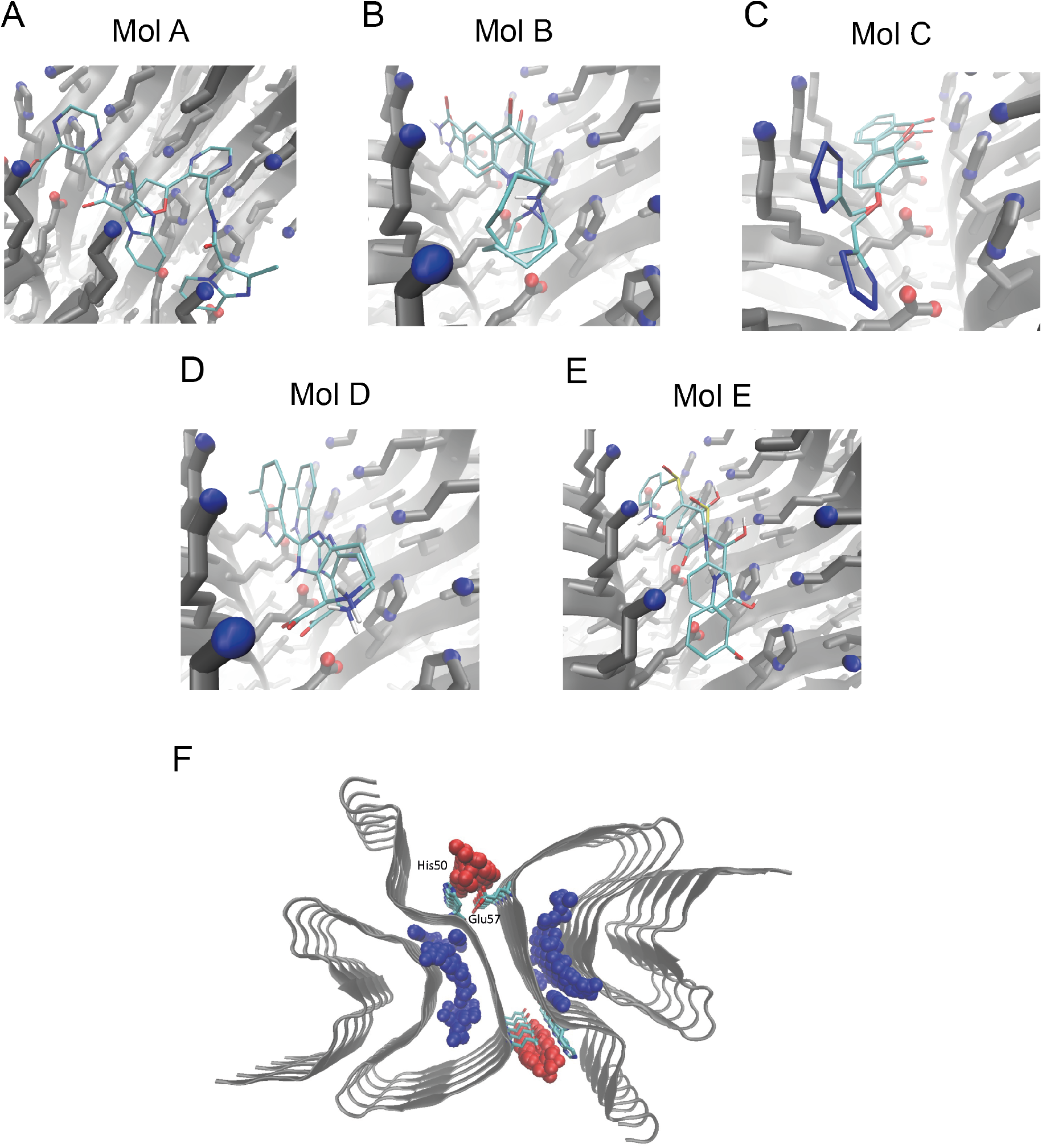
Computational docking of small molecules to α-synuclein fibrils. **(A-E)** Binding poses of small molecules A-E to α-synuclein fibrils binding pocket (centered between residues His50 and Glu57), determined either through FRED or AutoDock Vina. **(F)** Fibril structure (PDB:6cu7, cyan), pockets in sphere representation, with blue signifying pockets at the fibril core, and red at the fibril surface. Key binding site residues His50 and Glu57 are shown in licorice representation.

To rule out any potential effects whereby the small molecules inhibit the aggregation of α-synuclein by stabilising non-fibrillar aggregates, we used transmission electron microscopy (TEM) to image the α-synuclein species formed at the end of the aggregation reaction in the absence and presence of compound C (**Fig. S2**). These measurements showed a similar morphology of α-synuclein fibrils, suggesting that the inhibitors that we identified are able to delay the aggregation process, without redirecting it to affect the final amounts of α-synuclein aggregates formed (**Fig. S2**).

### Kinetic analysis of α-synuclein aggregation in the presence of the inhibitors

To characterise the inhibitory potency of the 5 positive compounds against the aggregation of α-synuclein, we measured the secondary nucleation process of α-synuclein in the presence of varying small molecule concentrations, from sub-stoichiometric ratios (0.625 molar equivalents) to over-stoichiometric ratios (5 molar equivalents) (**Figs. 4A** and **S3**). For all the 5 compounds, we observed a dose-dependent inhibition in the aggregation of α-synuclein, resulting in a systematic increase in the *t*_*1/2*_ of aggregation. Similarly to what we observed in the preliminary screening, the potency of the compounds varied between molecules A, B, and E, which exhibited a weaker effect, and molecules C and D, which had a stronger effect (**Figs. 3B** and **4A**). We further quantified the effect of the compounds, finding that they were able to significantly inhibit the rate of fibril amplification and change the fibril number concentration. We obtained these results by fitting the experimental data with a logistic function describing the amplification of α-synuclein aggregates over time (25) (Materials and Methods) (**Fig. 4B**). These compounds are likely to compete with α-synuclein monomers on the nucleation sites, as shown previously with other compounds and molecular chaperones that also exhibit inhibition of secondary nucleation processes (24, 26, 34, 35).

**Figure 4.**
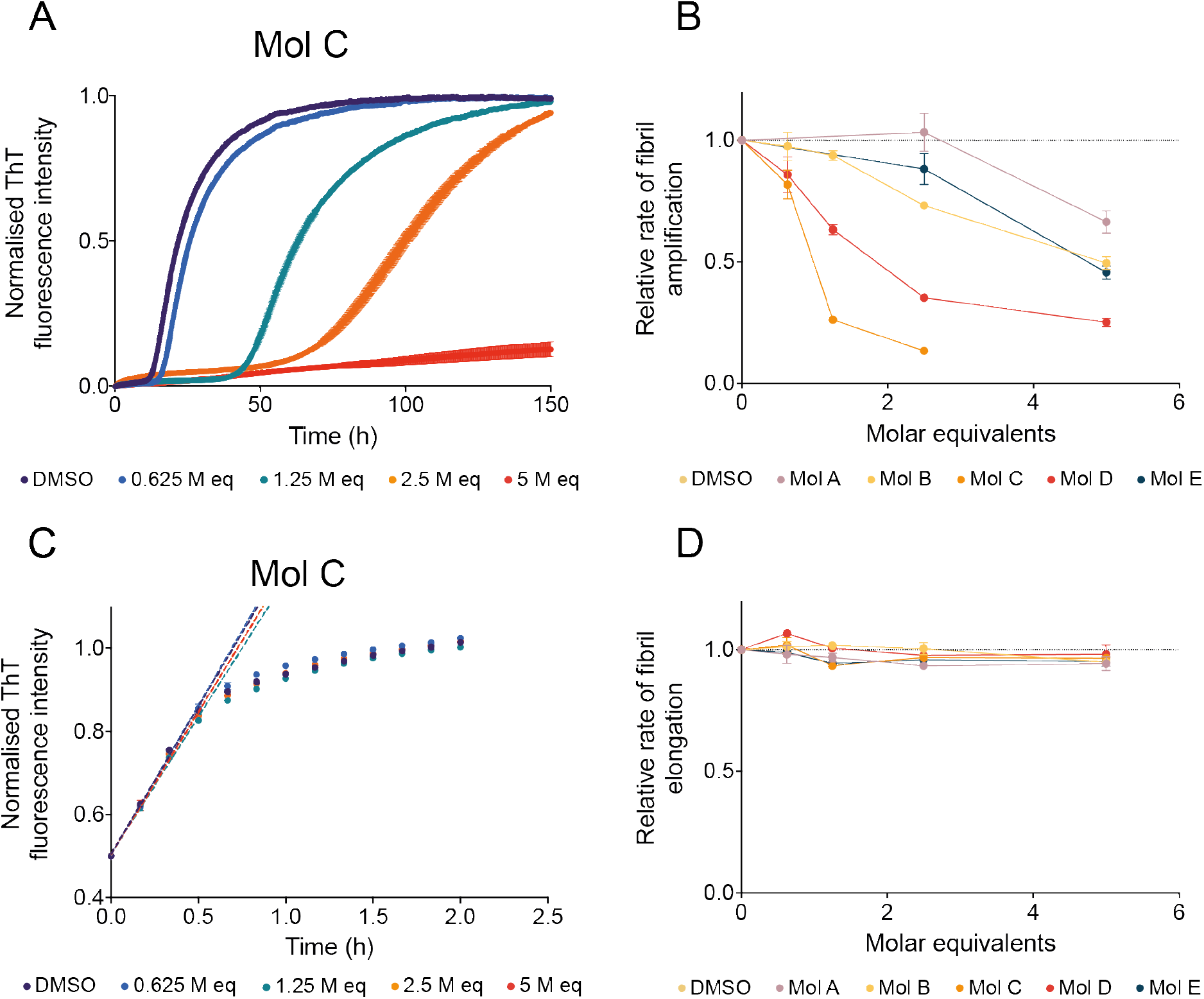
The small molecule identified by docking specifically inhibit the proliferation of α-synuclein aggregates by secondary nucleation. **(A)** Kinetic profiles of a 10 M solution of α-synuclein in the presence of 25 nM seeds at pH 4.8, 37 °C, either in the presence of 1% DMSO alone (purple), or of increasing molar equivalents of molecule C (represented in different colours). **(B)** Relative rate of fibril amplification of α-synuclein the presence of molecules A-E as shown in panel A and **Fig. S2**, normalised to the DMSO control. **(C)** Kinetic profiles of a 10 μM solution of α-synuclein in the presence of 5 M seeds at pH 4.8, 37 °C, either in the presence of 1% DMSO alone (purple), or in the presence of increasing molar equivalents of molecule C (represented in different colours). Dotted lines indicate the *v*_*max*_ of the reaction which is used to extract the elongation rate of the aggregation process. **(D)** The relative rate of fibril elongation of a-synuclein the presence of molecules A-E as shown in panel C and **Fig. S3**, normalised to the DMSO control.

The potency in inhibiting the amplification rate of the compounds was found to be in the order C > D > B and E > A. For instance, at 2.5 molar equivalents, while molecules C and D were able to inhibit the amplification rate of α-synuclein by 87% and 65%, respectively, compounds B and E were only able to inhibit this rate by 27% and 12%, respectively, and compound A was not able to significantly inhibit this process at this molar equivalent concentration.

To further probe the mechanism by which the compounds inhibit the aggregation process of α-synuclein, we also measured the aggregation process of α-synuclein in the absence and presence of the compounds with the addition of high concentration of preformed α-synuclein fibril seeds (**Figs. 4C** and **S4**). Under such conditions, the aggregation process of α-synuclein proceeds as an exponential rather than sigmoidal function, indicative of an aggregation mechanism dominated by elongation processes with negligible contribution of secondary nucleation (36, 37) (**Fig. 4C**). This phenomenon can be ascribed to the large number of growth-competent ends of fibrils present at the start of the aggregation process for further monomer addition. We observed that, under such conditions, the elongation rate, and consequently the aggregation process, was not significantly perturbed by the presence of the compounds (**Figs. 4C,D** and **S4**). Thus, the inhibition, as observed using the low seeded aggregation assay, is likely due to the inhibition of the secondary nucleation process rather than the elongation process (**Figs. 4A** and **S3**). This also indicates that the molecules that were identified using our approach are likely binding the surface of α-synuclein fibrils, rather than the ends, further supporting their binding to the selected groove.

### Molecule C inhibits α-synuclein oligomer formation

The reactive flux towards α-synuclein oligomers in the aggregation reaction of α-synuclein is governed by multiple processes, crucially secondary nucleation (29). By extracting the change in the amplification rate due to the presence of the small molecules, and by accounting for the specific inhibition of the secondary nucleation process rather than the elongation process, we can calculate the reactive flux towards oligomers over time in the absence and presence of increasing concentrations of the molecules (**Figs. 5A** and **S5**) (Materials and Methods). Depending on their potency, we observed that compounds were able to delay the rate of reactive flux towards oligomers, as well as a delay in the overall formation of oligomers over time (integral area of flux) (25, 28).

**Figure 5.**
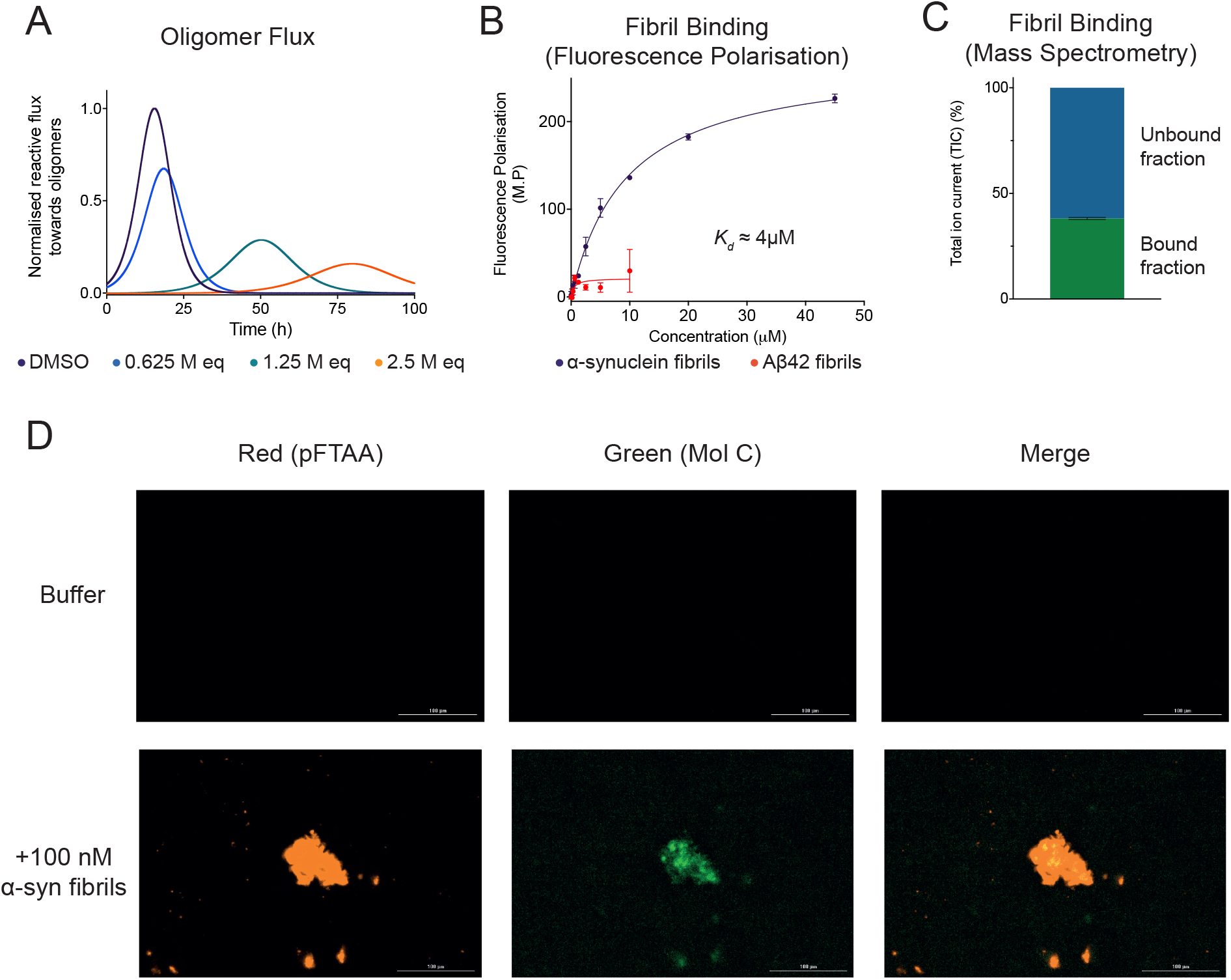
Molecule C inhibits the reactive flux towards α-synuclein oligomers and displays binding affinity and specificity towards α-synuclein fibrils. **(A)** Time dependence of the reactive flux towards α-synuclein oligomers either in the presence of 1% DMSO alone (purple) or in the presence of increasing molar equivalents of molecule C (represented in different colours), normalised to the DMSO control. **(B)** Change in fluorescence polarisation (in mP units) of 10 μM molecule C with increasing concentrations of either α-synuclein fibrils (purple) or A 42 fibrils (red). The solid lines are fits to the points using a one-step binding curve, estimating a *K*_*d*_ of 4 μM for molecule C towards α-synuclein fibrils. **(C)** Total ion current (TIC) of 10 μM molecule C bound and unbound to 10 μM α-synuclein fibrils detected by mass spectrometry (see Materials and Methods). **(D)** Representative images indicating either the fluorescence of the red channel (amyloid-specific dye pFTAA) or the green channel (molecule C) following incubation in the absence (top) or presence (bottom) of 100 nM α-synuclein fibrils.

### Molecule C binds α-synuclein fibrils

As a validation of the binding of these candidates to the surface of α-synuclein fibrils, we also performed fluorescence polarisation experiments of molecule C, the strongest inhibitor of all the positive compounds, in the presence of increasing concentrations of α-synuclein fibrils (**Figs. 5B** and **S6**). Additionally, since molecule C exhibits a high degree of intrinsic fluorescence, fluorescence polarisation measurements allowed us to determine the proportion of bound and unbound fractions of molecule C towards α-synuclein fibrils (**Fig. S6**). We observed a dose-dependent increase in the polarization (in millipolarization units, mP) of molecule C as a function of increasing concentrations of α-synuclein fibrils. By fitting this response as a function of the concentration of α-synuclein fibrils, the apparent dissociation constant (*K*_*d*_) was found to be about 4μM (**Fig. 5B**) (Materials and Methods). However, we note that the *K*_*d*_ value obtained is based on the concentration of α-synuclein fibrils in monomer equivalents. Since α-synuclein fibrils tend to contain at least 200 monomers per unit (36), it is thus likely that the actual *K*_*d*_ value (in terms of number of fibrillar units) should be significantly lower.

To test for the specificity of molecule C for α-synuclein fibrils, we measured the change in polarization upon incubation of molecule C with A β42 fibrils, which are associated with Alzheimer’s disease (**Fig. 5B**). In this case, we observed a much lower increase in the polarization, as only a slight increase could be measured at 10 μM of Aβ42 fibrils. This result suggests that molecule C binds specifically to the surface of α-synuclein fibrils, as predicted from the docking, rather than having generic non-specific interactions with hydrophobic aggregates (**Fig. 5B**).

To further support the fluorescence polarisation data, we also performed a mass-spectrometry based pull down assay to assess the amounts of molecules bound to α-synuclein fibrils (**Fig. 5C**) (Materials and Methods). We found that approximately 40% of molecule C was still associated to α-synuclein fibrils after an ultracentrifugation pull-down (**Fig. 5C**). Finally, images obtained from α-synuclein fibrillar samples containing molecule C or the amyloid-specific fluorescence dye pFTAA demonstrated the colocalisation of molecule C with the fibrillar aggregates (imaged by pFTAA) (**Fig. 5D**).

This behaviour was also found to be visualised in the presence of neuroblastoma cells, thus confirming the significant affinity of molecule C towards α-synuclein fibrils, as well as demonstrating the potential use of such molecules in biological studies involving α-synuclein aggregation (**Fig. S7**).

## Conclusions

In this work, we have demonstrated the use of an approach to identify and characterise small molecules whose mechanism of action is to bind α-synuclein fibrils and inhibit the secondary nucleation step in the autocatalytic proliferation of α-synuclein aggregates. We anticipate that such approach will enable the rational design and systematic development of small molecules capable of binding specific sites with different properties on α-synuclein fibrils of different morphologies, as well as in fibrils formed by other disease-related proteins, thereby creating new opportunities in the diagnostics and therapeutics for protein misfolding diseases.

## Materials and Methods

### Computational docking

The docking protocol used in this study comprises four stages. First, we determined a binding site on the α-synuclein fibrils. To achieve this goal, we analysed a structure of α-synuclein fibrils (PDB ID: 6cu7) using Fpocket (38), which identifies small molecule pockets based on volume criteria (**Fig. S1**). The resulting pockets were categorised in: (a) pockets in the fibril core surrounded by His50 to Lys58 and Thr72 to Val77, and (b) pockets on the fibrils surface, surrounded by His50 and Glu57. Since α-synuclein secondary nucleation has been reported as significant only below pH 5.8 (36), when histidine is protonated, we further reduced the space of possible binding pockets, selecting the binding site containing His50 and Glu57.

For the selection of screening compounds we used the ZINC library (32). In order to reduce the chemical space of small molecules considered in the docking calculations, central nervous system multiparameter optimization (CNS MPO) criteria (39) were applied, effectively reducing the space to approximately 2 million compounds. We subjected these compounds to docking calculation against the binding site identified above using AutoDock Vina (30). To increase the confidence of the calculations, the top-scoring 10,000 small molecules were selected and docked against the same α-synuclein binding site, using FRED (OpenEye Scientific Software) (31). The top-scoring, common 1,000 compounds in both docking protocols are selected and clustered using Tanimoto clustering (40), leading to a list of 79 clusters.

### Preparation of compounds and chemicals

The centroids from the above 79 clusters were selected for experimental validation. Compounds were purchased from MolPort (Riga, Latvia), and in the cases for which centroids were not available for purchase, the molecules in the clusters with the closest chemical structures were used as the representative compounds instead. In the end, a total of 67 compounds were purchased (centroids and alternative compounds in 12 clusters were all not available for purchase), and then prepared in DMSO to a stock of 5 mM. All chemicals used were purchased at the highest purity available.

### Preparation of α-synuclein

Recombinant α-synuclein was purified as described previously (36, 37, 41). The plasmid pT7-7 encoding for human α-synuclein was transformed into BL21-competent cells. Following transformation, competent cells were grown in lysogeny broth (LB) in the presence of ampicillin (100 μg/mL). Cells were induced with isopropyl β-D-1-thiogalactopyranoside (IPTG), and grown overnight at 37°C for 4 h and harvested by centrifugation in a Beckman Avanti J25 centrifuge with a JA-20 rotor at 5,000 rpm (Beckman Coulter, Fullerton, CA). The cell pellet was resuspended in 10 mM Tris, pH 8.0, 1 mM EDTA, 1 mM PMSF and lysed by multiple freeze–thaw cycles and sonication. The cell suspension was boiled for 20 min and centrifuged at 13,500 rpm with a JA-20 rotor (Beckman Coulter). Streptomycin sulfate was added to the supernatant to a final concentration of 10 mg/mL and the mixture was stirred for 15 min at 4 °C. After centrifugation at 13,500 rpm, the supernatant was taken with an addition of 0.36 g/mL ammonium sulfate. The solution was stirred for 30 min at 4 °C and centrifuged again at 13,500 rpm. The pellet was resuspended in 25 mM Tris, pH 7.7, and ion-exchange chromatography was performed using a HQ/M-column of buffer A (25 mM Tris, pH 7.7) and buffer B (25 mM Tris, pH 7.7, 600 mM NaCl). The fractions containing α-synuclein (≈ 300 mM) were dialysed overnight against the appropriate buffer. The protein concentration was determined spectrophotometrically using the extinction coefficient ε280 = 5600 M^-1^ cm^-1^.

### Preparation of α-synuclein fibril seeds

α-synuclein fibril seeds were produced as described previously (36, 37). Samples of α-synuclein (700 μM) were incubated in 20 mM phosphate buffer (pH 6.5) for 72 h at 40 °C and stirred at 1,500 rpm with a Teflon bar on an RCT Basic Heat Plate (IKA, Staufen, Germany). Fibrils were then diluted to 200 μM, aliquoted and flash frozen in liquid nitrogen, and finally stored at −80 °C. For the use of kinetic experiments, the 200 μM fibril stock was thawed, and sonicated for 15 s using a tip sonicator (Bandelin, Sonopuls HD 2070, Berlin, Germany), using 10% maximum power and a 50% cycle.

### Kinetic assays

α-Synuclein was injected into a Superdex 75 10/300 GL column (GE Healthcare) at a flow rate of 0.5 mL/min and eluted in 20 mM sodium phosphate buffer (pH 4.8) supplemented with 1 mM EDTA. The obtained monomer was diluted in buffer to a desired concentration, and supplemented with 50 μM ThT and preformed α-synuclein fibril seeds. The molecules (or DMSO alone) were then added at the desired concentration to a final DMSO concentration of 1% (v/v). Samples were prepared in low-binding Eppendorf tubes, and then pipetted into a 96-well half-area, black/clear flat bottom polystyrene NBS microplate (Corning 3881), 150 μL per well. The assay was then initiated by placing the microplate at 37 °C under quiescent conditions in a platereader (FLUOstar Omega, BMG Labtech, Aylesbury, UK). The ThT fluorescence was measured through the bottom of the plate with a 440 nm excitation filter and a 480 nm emission filter.

### Transmission electron microscopy

10 μM α-synuclein samples were prepared and aggregated as described in the kinetic assay, in the absence or presence of 25 μM molecule C, without the addition of ThT. Samples were collected from the microplate at the end of the reaction (150 hours) into low-binding Eppendorf tubes. They were then prepared on 400-mesh, 3-mm copper grid carbon support film (EM Resolutions Ltd.) and stained with 2% uranyl acetate (wt/vol). The samples were imaged on an FEI Tecnai G_2_transmission electron microscope (Cambridge Advanced Imaging Centre). Images were analysed using the SIS Megaview II Image Capture system (Olympus).

### Determination of the elongation rate constant

In the presence of high concentrations of seeds (≈ μM), the aggregation of α-synuclein is dominated by the elongation of the added seeds (36, 37). Under these where other microscopic processes are negligible, the aggregation kinetics for α-synuclein can be described by (36, 37)

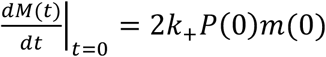

where *M(t)* is the fibril mass concentration at time *t, P(0)* is the initial number of fibrils, *m(0)* is the initial monomer concentration, and *k*_*+*_ is the rate of fibril elongation. In this case, by fitting a line to the early time points of the aggregation reaction as observed by ThT kinetics, 2*k*_*+*_*P(0)m(0)* can be calculated for α-synuclein in the absence and presence of the compounds. Subsequently, the elongation rate in the presence of compounds can be expressed as a normalised reduction as compared to the elongation rate in the absence of compounds (1% DMSO).

### Determination of the fibril amplification rate constant

In the presence of low concentrations of seeds, the fibril mass fraction *M(t)* over time was described using a generalised logistic function to the normalised aggregation data (25, 42)

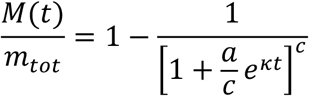

where *m*_*tot*_ denotes the total concentration of α-synuclein monomers. The parameters a and c are defined as

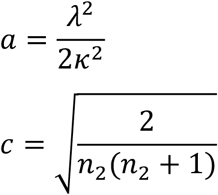

The parameters λ and κ represent combinations for the effective rate constants for primary and secondary nucleation, respectively, and are defined as (25, 42):

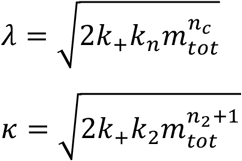

where *k*_*n*_ and *k*_*2*_ denote the rate constants for primary and secondary nucleation, respectively, and *n*_*c*_ and *n*_*2*_ denote the reaction orders of primary and secondary nucleation, respectively. In this case, c was fixed at 0.3 for the fitting of all data, and *k*_*2*_, the amplification rate, is expressed as a normalised reduction for α-synuclein in the presence of the compounds as compared to in its absence (1% DMSO).

### Determination of the oligomer flux

The prediction of the reactive flux towards oligomers over time was calculated as (25, 42)

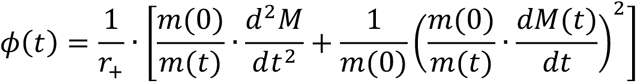

where *r*_*+*_ *= 2k*_*+*_*m(0)* is the apparent elongation rate constant extracted as described earlier, and *m(0)* refers to the total concentration of monomers at the start of the reaction.

### Fluorescence polarisation

10 μM of molecule C was incubated with increasing concentrations of either pre-formed α-synuclein or Aβ42 fibrils (in 1% DMSO). After incubation, the samples were pipetted into a 96-well half-area, black/clear flat bottom polystyrene nonbinding surface (NBS) microplate (Corning 3881). The fluorescence polarisation of molecule C was monitored using a plate reader (CLARIOstar, BMG Labtech, Aylesbury, UK) under quiescent conditions at room temperature, using a 360 nm excitation filter and a 430 emission filter.

### Mass spectrometry

10 μM of preformed α-synuclein fibrils were incubated with 10 μM of molecule C in 20 mM sodium phosphate buffer (pH 4.8) supplemented with 1 μmM EDTA overnight under quiescent conditions at room temperature. The samples were then ultracentrifuged at 100,000 *g* for 30 min and the supernatant was removed for analysis using a Waters Xevo G2-S QTOF spectrometer (Waters Corporation, MA, USA).

### Cell cultures

Human SH-SY5Y neuroblastoma cells (A.T.C.C., Manassas, VA) were cultured in Dulbecco’s modified Eagle medium (DMEM)-F12+GlutaMax supplement (Thermo Fisher Scientific, Waltham, MA) with 10% heat-inactivated fetal bovine serum. The cell cultures were maintained in a 5.0% CO2 humidified atmosphere at 37 °C and grown until 80% confluence for a maximum of 20 passages.

### Colocalisation assay

The cells were plated into a 96-well plate and treated for 24 h with samples containing α-synuclein fibrils at 100 μnM concentration (monomer equivalents) pre-incubated with pFTAA, molecule C, or buffer (20 mM sodium phosphate buffer, pH 4.8). After treatment, the cells were stained with Amytracker 630 (Ebba Biotech AB, Sweden). Images were acquired using the fluorescence microscope Cytation5 Cell Imaging Reader (BioTek Instruments, Winooski, VT).

## Supplementary Information

**Figure S1.**
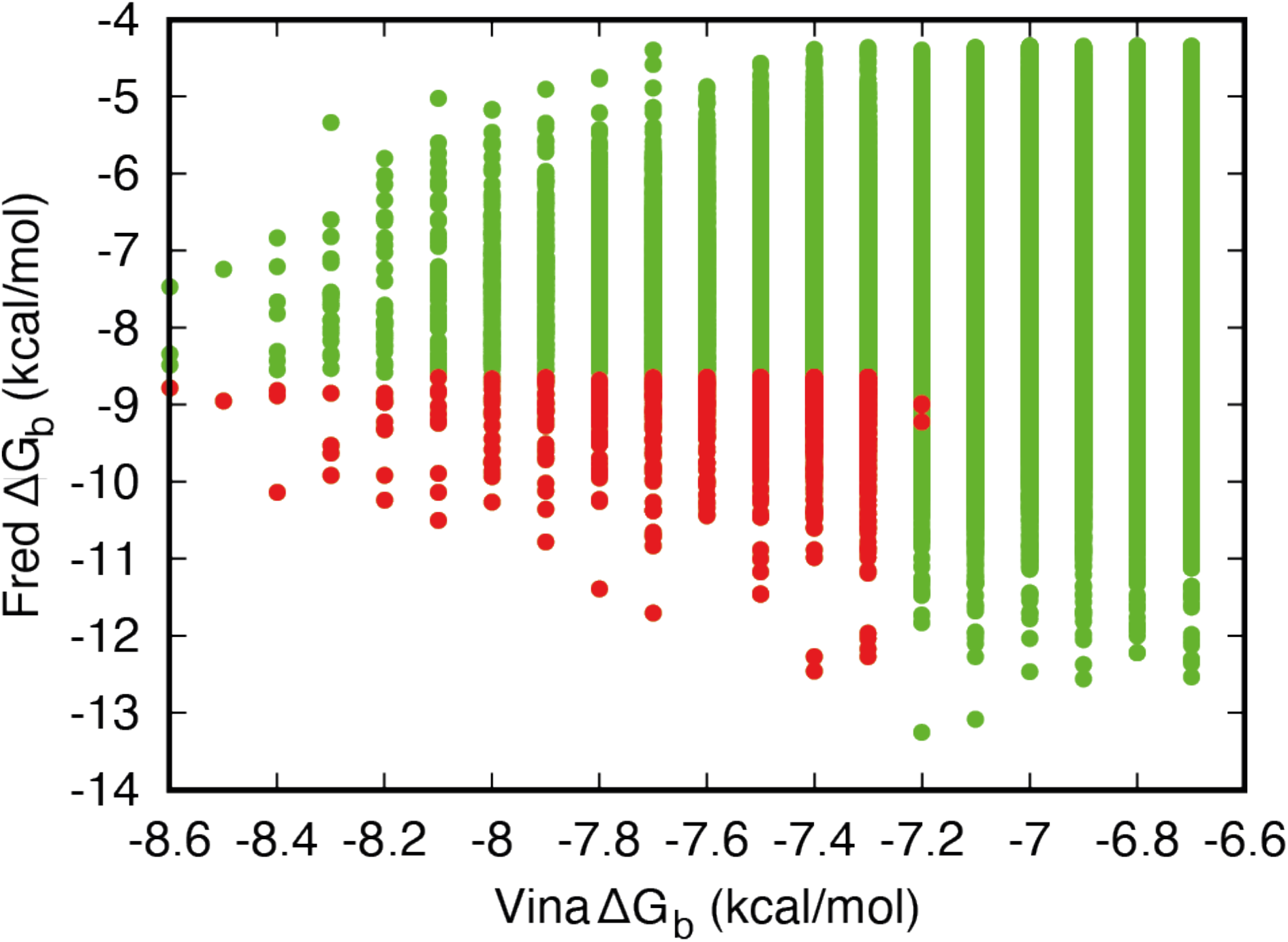
Distribution of the predicted binding affinity for α-synuclein fibrils of the top 10,000 compounds screened via computational docking. The predicted binding score (ΔG_b_) of each compound calculated either via FRED (Fred) or AutoDock Vina (Vina). Points in red denote the top 1,000 compounds with the highest predicted binding scores calculated by both methods, which were further selected as an enriched compound library of potential α-synuclein fibril binders.

**Figure S2.**
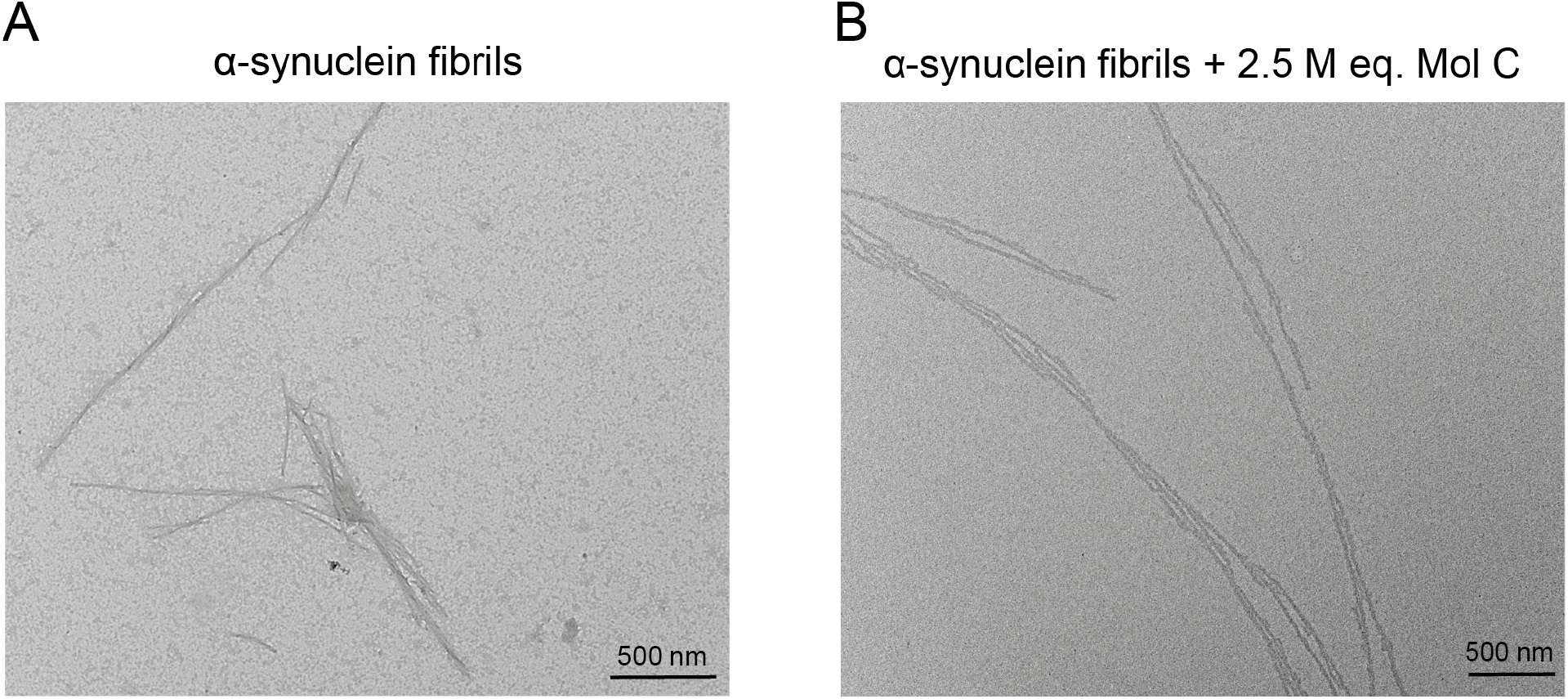
Comparison of TEM images of α-synuclein fibrils generated in the absence and presence of the small molecule inhibitor C. **(A,B)** TEM images of 10 μM α-synuclein fibrils grown in the absence (**A**) or presence (**B**) of 2.5 molar equivalents of molecule C.

**Figure S3.**
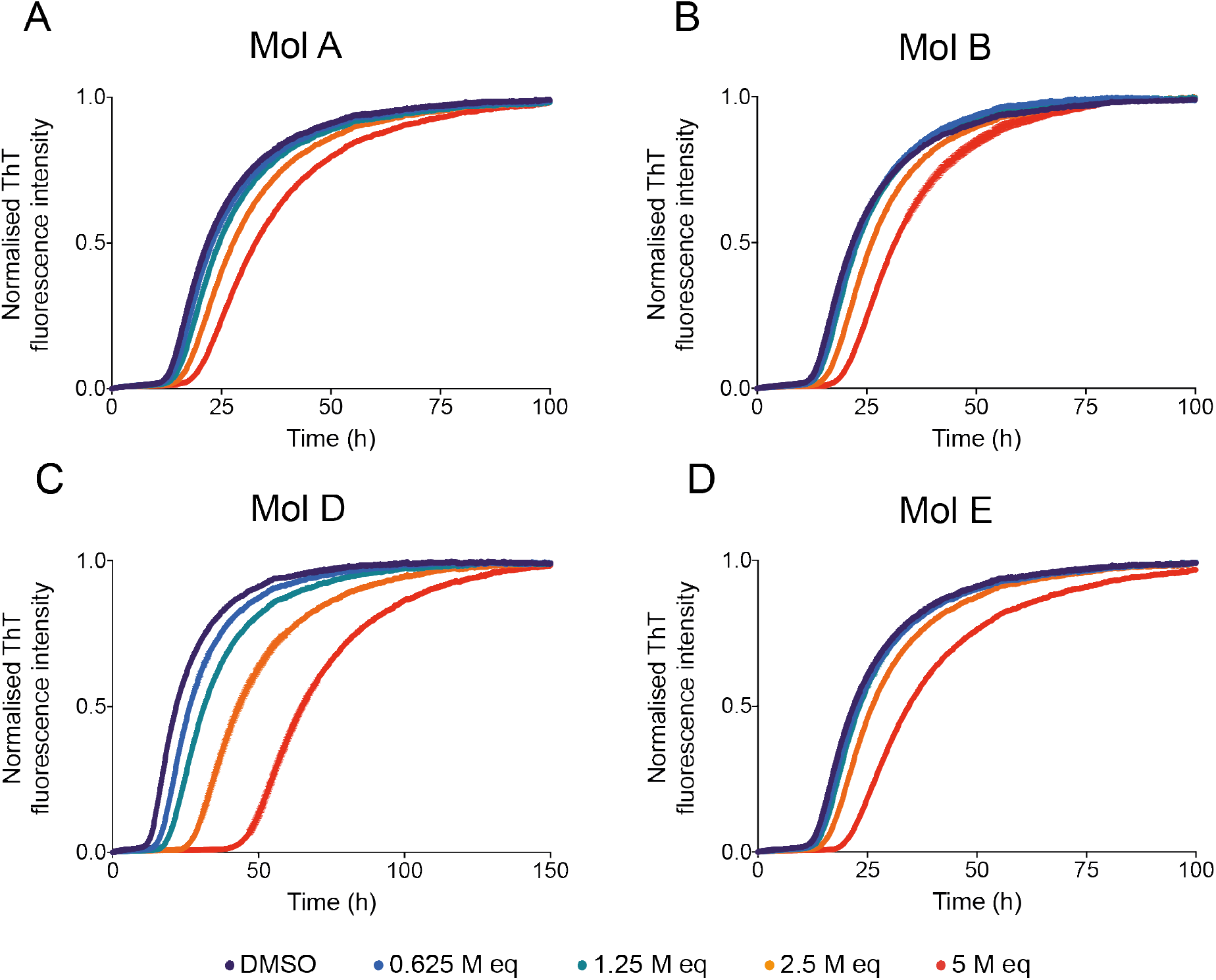
Positive compounds from the docking library inhibit the surface-catalysed secondary nucleation process of α-synuclein aggregation. **(A-D)** Kinetic profiles of a 10 μM solution of α-synuclein in the presence of 25 nM seeds at pH 4.8, 37 °C, either in the presence of 1% DMSO alone (purple), or in the presence of increasing molar equivalents of either molecule A (**A**), molecule B (**B**), molecule D (**C**), or molecule E (**D**).

**Figure S4.**
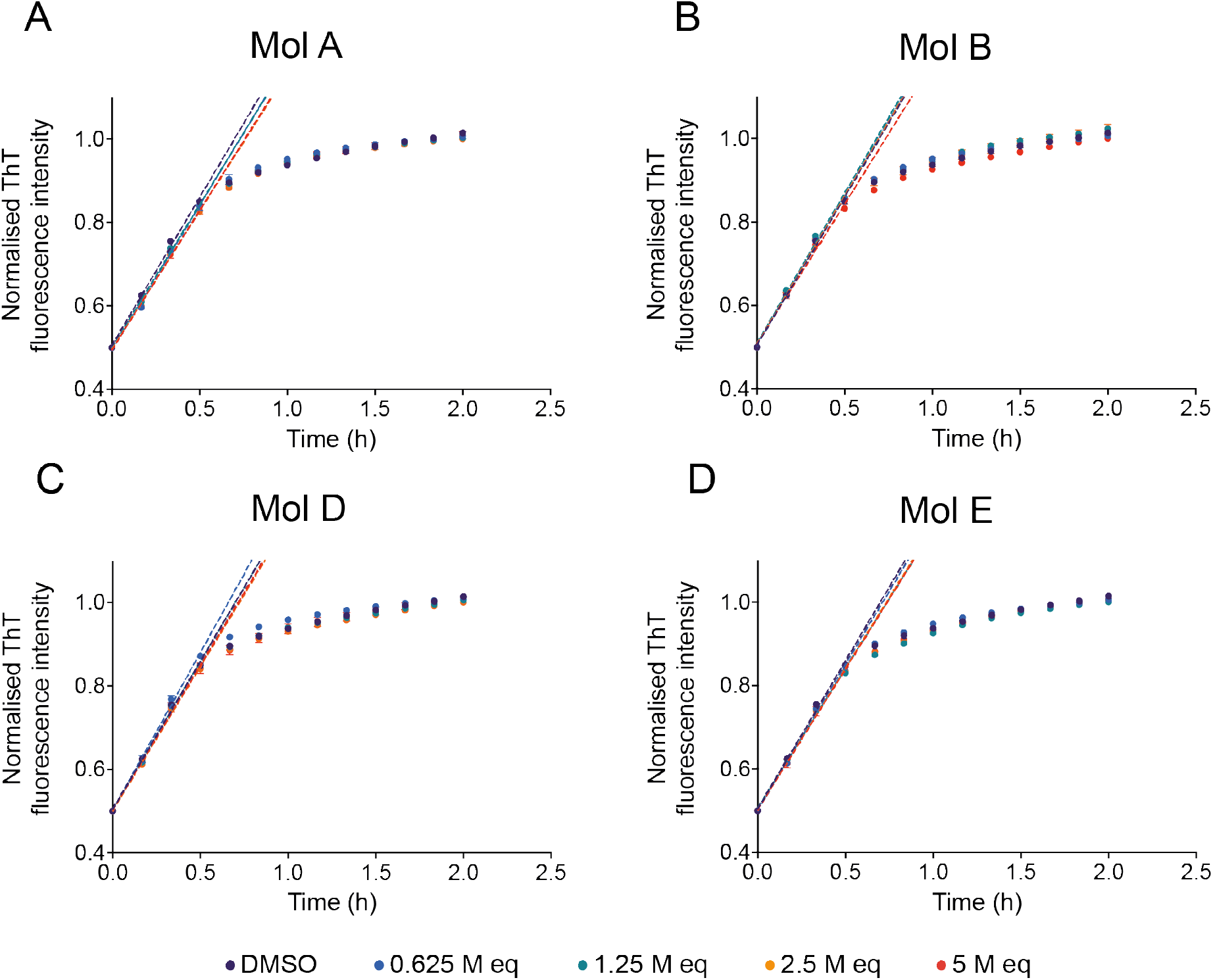
Positive compounds from the docking library do not significantly inhibit the elongation process of α-synuclein aggregation. **(A-D)** Kinetic profiles of a 10 μM solution of α-synuclein in the presence of 5 μM seeds at pH 4.8, 37°C, either in the presence of 1% DMSO alone (purple), or in the presence of increasing molar equivalents of either molecule A (**A**), molecule B (**B**), molecule D (**C**), or molecule E (**D**). Dotted lines indicate the *v*_*max*_ of the reaction which is used to extract the elongation rate of the aggregation process (**Fig. 4D**).

**Figure S5.**
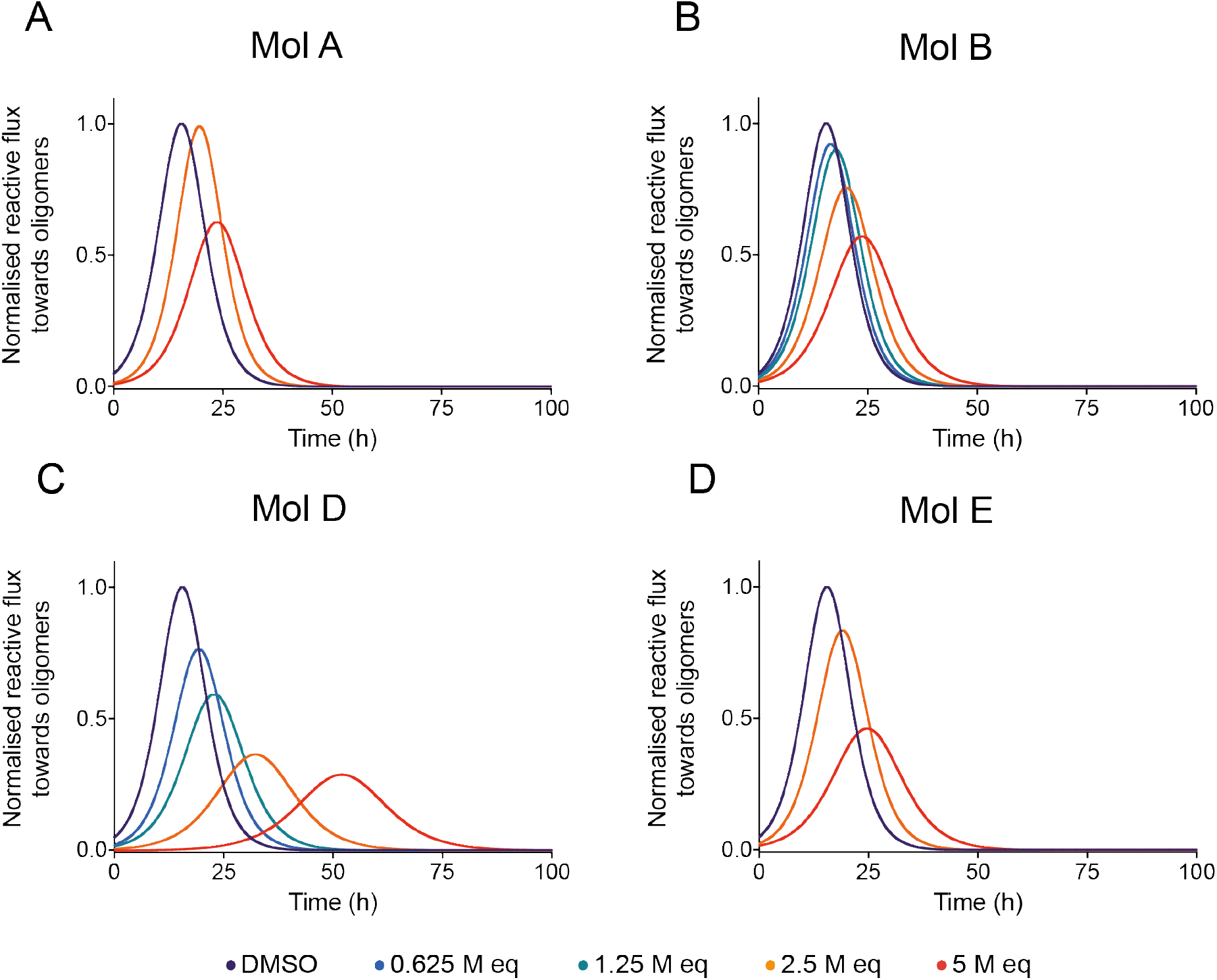
Positive compounds from the docking library inhibit the reactive flux towards α-synuclein oligomers. **(A-D)** Time dependence of the reactive flux towards α-synuclein oligomers either in the presence of 1% DMSO alone (purple) or in the presence of increasing molar equivalents of molecule A **(A)**, molecule B (**B**), molecule D (**C**), or molecule E (**D**) normalised to the DMSO control.

**Figure S6.**
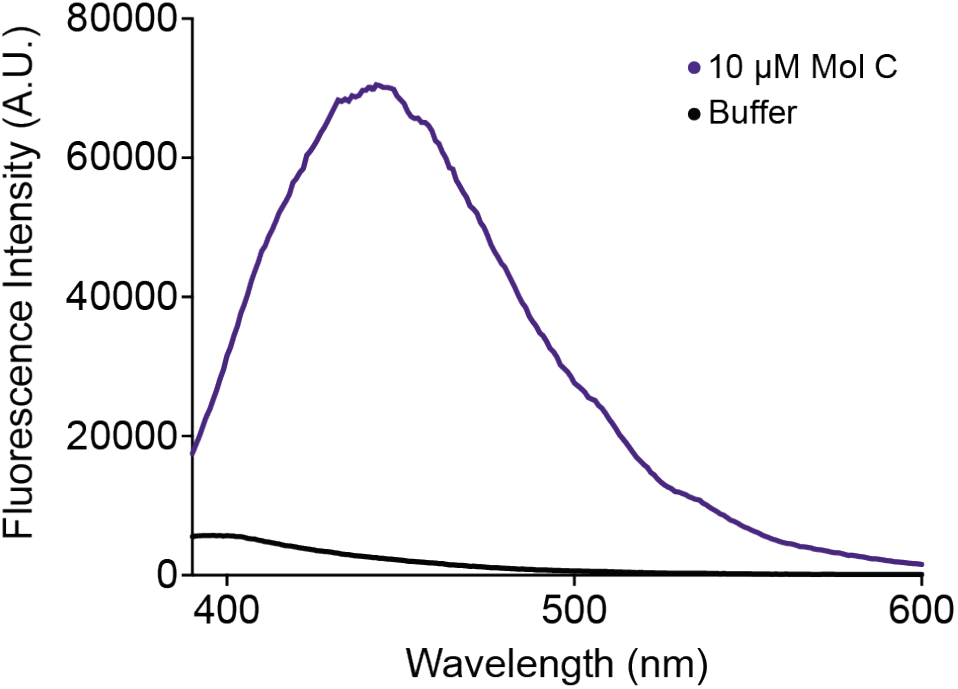
Fluorescence emission spectra of molecule C. Fluorescence emission spectra of either buffer (black) or 10 μM molecule C (purple) in sodium phosphate buffer, pH 4.8, 1 mM EDTA (λ ex = 360 nm).

**Figure S7.**
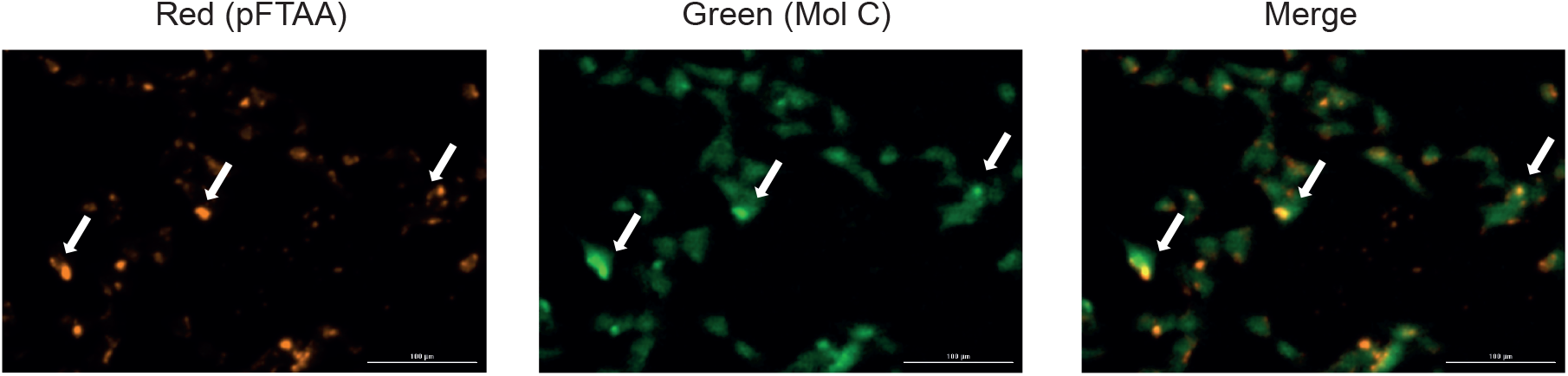
Co-localisation of molecule C with α-synuclein fibrils in the presence of neuroblastoma cells. Representative images indicating either the fluorescence of the red channel (amyloid-specific dye pFTAA) or the green channel (molecule C) following incubation in the presence of 100 nM α-synuclein fibrils and neuroblastoma cells. White arrows indicate the specific colocalisation of molecule C with α-synuclein fibrils in the presence of cells.

## Notes

### Competing Interest Statement

The authors have declared no competing interest.

